# Resolving Sialylated N-Glycans and Immune Cell Landscapes Using a Unified Same-Section IMC–MSI Workflow

**DOI:** 10.64898/2026.02.21.707206

**Authors:** Jake Griner, Reto Gerber, Mark D. Robinson, Carsten Krieg, Silvia Guglietta

## Abstract

Integrating antibody-based imaging with mass spectrometry imaging (MSI) on the same formalin-fixed paraffin-embedded (FFPE) tissue section offers powerful opportunities for multimodal spatial analysis but remains analytically challenging due to cross-platform chemical and physical interference. In particular, chemically aggressive on-tissue derivatization strategies required for isomer-resolved glycan MSI may compromise downstream antibody detection.

Here, we systematically evaluate the analytical compatibility and acquisition order of imaging mass cytometry (IMC) and sialic-acid-linkage-resolving N-glycan MALDI-MSI using an Amidation–Activation–X-Linkage (AAXL) derivatization strategy on the same FFPE tissue section. Two same-section workflows were compared: AAXL–MALDI MSI followed by IMC (MALDI-first) and IMC followed by AAXL–MALDI MSI (IMC-first). We find that AAXL-first processing results in severe and widespread loss of IMC antibody signal across epithelial, immune, and nuclear markers, rendering subsequent antibody-based analysis unreliable. In contrast, IMC-first acquisition preserves quantitative antibody performance while maintaining spatial glycan distributions, relative abundance structure, and isomer-specific signal integrity in downstream AAXL–MALDI MSI.

Using high-precision co-registration, we further demonstrate that IMC-first sequencing enables analytically robust integration of IMC and MSI data at both domain and pixel levels. These results establish IMC-first acquisition as the preferred same-section strategy for workflows combining antibody imaging with chemically intensive, isomer-resolved glycan MSI and provide generalizable guidance for the design of multimodal spatial mass spectrometry experiments.

## Introduction

In recent years, highly multiplexed imaging techniques have revolutionized the field of spatial biology. Antibody-based multiplexed imaging modalities resolve cell identity and functional states at single-cell and subcellular resolution through predefined protein targets, whereas non–antibody-based approaches such as matrix-assisted laser desorption/ionization mass spectrometry imaging (MALDI-MSI) provide unbiased spatial profiling of glycans, lipids, and metabolites without a priori probes [1]. These approaches are analytically complementary, motivating increasing efforts to integrate them into unified spatial workflows.

Despite this complementarity, combining antibody-based imaging and MALDI-MSI on the same FFPE tissue section remains analytically challenging. Antibody-based multiplexed imaging modalities such as cyclic IF [2], CODEX [3], MIBI [4] and imaging mass cytometry [5] resolve cell types at single-cell and in many cases even sub-cellular resolution [6]. Most commercially available MALDI-MSI instruments operate at spatial resolutions of 10 µm to 30 µm [7], while improved resolution MALDIMSI is an area of active research, with some custom instruments achieving 1 µm resolution [8]. The IMC to MALDI-MSI resolution mismatch complicates precise data co-registration. Although adjacent-section strategies partially mitigate this issue, we and others have shown that even micrometer-scale misalignment introduces substantial analytical error, making same-section multimodal acquisition preferable [9]. However, same-section integration raises critical concerns regarding chemical compatibility, acquisition order, and preservation of both antibody epitopes and analyte integrity.

Prior same-section studies have demonstrated feasibility under restricted conditions but they fall into two distinct categories that matter for experimental design: (i) **lipid** MALDI-MSI combined with antibody imaging, and (ii) **glycan** MALDI-MSI combined with antibody imaging; and, separately, (a) **immunofluorescence (IF)**–based antibody methods (e.g., CODEX, cyclic IF) versus (b) **IMC** (metal-tag, laser ablation). Specifically, lipid MALDI-MSI performed *before* antibody imaging has shown minimal impact on downstream staining when paired with IF [10] or with IMC [11]. Likewise, N-glycan MALDI-MSI followed by IF (CODEX) on formalin fixed paraffin embedded (FFPE) tissues has been reported with little apparent loss of staining quality [12]. We also previously observed only mild to moderate disruptions of antibody staining performance when performing N-glycan MALDI-MSI prior to IMC [13]. However, these studies did not employ chemically aggressive on-tissue derivatization strategies required to resolve sialic acid linkage isomers, which are analytically and biologically important but highly labile during MALDI ionization.

Sialyation is a form of post-translational modification occurring on glycoproteins and glycolipids, which plays a key role in cellular communication and functions. In addition to their role in preventing cell adhesion to the endothelium and mediating intracellular signaling by interacting with sialic acid-binding immunoglobulin-like lectins (SIGLECs) and selectins, sialic acids participate in immune responses by regulating leucocyte trafficking and immune cell maturation and activation [14–16]. As such, the sialyation process is involved in multiple pathologies ranging from cancer to neurologic disorders, which explains the growing interest in the development of sialic acid-based therapeutics [7].

Analytically, MALDI MSI of sialic acids is challenging due to fragmentation during ionization [17]. Stabilization and differentiation of *α*2,3- and *α*2,6 -linked sialic acids provides a critical analytical advantage, as these terminal glycans regulate immune recognition, receptor signaling, cell–cell interactions, and disease-associated tissue remodeling. The Amidation–Activation–X-Linkage (AAXL) strategy enables in situ stabilization and mass differentiation of these linkage isomers but requires sequential carbodiimide activation, nucleophilic amidation, and high-temperature organic incubations [18]. Whether such chemically intensive workflows are compatible with downstream antibody-based imaging—particularly imaging mass cytometry (IMC)—has not been systematically evaluated.

Here, we present the first direct analytical assessment of same-section compatibility between AAXL-enabled N-glycan MALDI-MSI and IMC. We performed a controlled comparison of two acquisition orders:(i) AAXL–MALDI-MSI followed by IMC and (ii) IMC followed by AAXL–MALDI-MSI. We demonstrate that AAXL-first processing severely compromises IMC antibody staining, whereas IMC-first acquisition preserves both antibody performance and glycan spatial integrity. These findings establish IMC-first acquisition as the preferred strategy for same-section workflows requiring sialic acid linkage–resolved spatial glycomics and define practical compatibility limits for chemically intensive multimodal imaging.

## Experimental Procedure

### Human Tissue Collection

This study utilized formalin fixed paraffin embedded (FFPE) liver specimens obtained from patients receiving standard of care treatment at the Medical University of South Carolina (IRB protocol #111306). Representative 1.5mm cores were compiled into a tissue microarray (TMA) by the Biorepository and Tissue Analysis (BTA) Shared Resource at the Hollings Cancer Center. All specimens were de-identified, assigned unique numeric codes, and securely stored to maintain patient confidentiality. Demographic data are provided in Supplementary Table S1.

### Imaging Mass Cytometry

#### Antibody panel design, conjugation, and validation

We assembled a metal-tagged antibody panel to capture liver architecture and inflammation, including epithelial, endothelial, and biliary markers, as well as immune lineages. Antibody clones were selected based on prior literature and empirical performance. Prior to this study, titrations were performed on tissue microarrays (Biomax) spanning diverse positive/negative control and final working concentrations were chosen to maximize signal-to-background separation. Full details including research resource identifiers (RRIDs) are listed in Supplementary Table S2.

#### IMC staining and acquisition

Briefly, 5 µm sections were deparaffinized, subjected to heat-induced epitope retrieval (Tris–EDTA, pH 9), and blocked (TBS,Triton X-100,BSA). Slides were incubated overnight at 4 ^°^C with the antibody cocktail, followed by next day washing and DNA labeling with intercalator-Ir. Regions of interest ( ROI, ∼,300 µm× 1,300 µm) were acquired on the Helios/Hyperion Tissue Imager (Standard BioTools). Instrument performance was verified with the 3-Element tuning slide, targeting ^175^Lu counts >1,000. Images were collected at 1 µm step size (200 Hz, −1 dBa) and visualized in MCD Viewer (v1.0.560.6). A full, step-by-step protocol with reagent compositions, wash sequences, and timing is provided in the Supplementary Methods.

### MALDI N-Glycan Imaging

#### Sialic acid stabilization

FFPE sections were dewaxed (xylenes) and rehydrated (ethanol gradients to HPLC-grade water), then briefly desiccated under vacuum. Differential stabilization of *α*2, 3 vs. *α*2, 6 sialic acids followed the AAXL amidation strategy [18]. In short, an on-tissue activation mix (EDC/HOBt with dimethylamine in DMSO) was applied and incubated at 60 ^°^C for 1 h to derivatize *α*2, 3 sialic acids (+37.03 Da). Slides were rinsed with DMSO and aspirated under vacuum. A second on-tissue reaction (propargylamine in DMSO; 60 ^°^C, 2 h) stabilized *α*2, 6 linkages (+27.05 Da), followed by rapid organic rinses (ethanol, Carnoy’s) and a brief 0.1% TFA/ethanol wash. Full reagent compositions, volumes, and wash timings are detailed in the Supplementary Methods.

#### MALDI MSI slide preparation

Heat-induced retrieval used 10 mM citraconic anhydride buffer (pH 3). PNGase F (PNGase F Prime, N-zyme Scientifics) was applied by automated spraying (M5, HTX Technologies), followed by enzymatic release glycan in a preheated humidity chamber (37.5 ^°^C, 2 h). CHCA matrix (7 mg/mL in 50% acetonitrile/0.1% TFA) was then sprayed (M5, HTX Technologies). Additional setup details and pass-by-pass parameters are provided in the Supplementary Methods.

#### MALDI-QTOF MSI acquisition

Imaging was performed on a timsTOF fleX MALDI-QTOF (Bruker Daltonics) operated in qTOF mode with TIMS disabled; the SmartBeam 3D laser was run at 10 kHz. External mass calibration used ESI-L Tune Mix (Agilent). Global instrument settings are summarized in Supplementary Table S3; regions of interest were approximately 1,600 µm × 1,600 µm.

### Statistical Analysis

#### IMC processing and cell annotation

Processing of IMC data to obtain single cells and cell types via MIMIC [9]. In short, raw IMC data was converted to multi-channel tiff images using steinbock [19] with additionally removing hot pixels and denoising (setting pixel values lower than 1.5 to zero). This was followed by segmentation using Mesmer [20], mean intensity aggregation and asinh transformation (with a cofactor of 1). Filtering of cells was done based on the following criteria: cell area smaller than 10 pixels, cells touching the border of an image, no cell marker signal. Integration of cells was done using harmony [21], followed by clustering using Leiden [22] and manual annotation of clusters.

#### Domain detection

The two domains (regenerative hepatocytes, bridging fibrotic bands) were determined based on the intensity of channels HepPar1 and Collagen-1, respectively. First, each of the two channels was normalized by the 99th quantile followed by assigning each cell to the domain with the higher intensity of its corresponding channel. Next, for each cell the mean intensity of its neighbors was calculated and the above scaling and classification was repeated, resulting in a smoothing of the domains resulting in more realistic borders.

#### MSI processing

A list of possible, known glycan m/z values was the starting point for the generation of a peak list. For each MSI pixel, the signal to noise ratio (SNR) was calculated using Cardinal [23] with the method “diff”. All available peaks were then filtered to only include peaks where at least 0.5% of all pixels had a SNR larger than 3. Raw spectra were first divided by the total ion current (TIC) followed by alignment to the reference peak list. The peaks were binned and the maximum signal was used for downstream processing. Finally, the extracted m/z values were log2 transformed.

#### Coregistration of MSI and IMC

The coregistration of MSI and IMC was conducted using MIMIC [9]. In short, intermediate microscopy slide scans were used as a scaffold for coregistration. The coregistration of MSI to the slide scan after MSI acquisition was based on detecting the physical location of ablation marks from the laser. The IMC coregistration to the “after IMC” acquisition slide scan was done similarly by detecting the physical ablation area. Stringent quality control regarding the precision of registration was done and samples with low precision were removed from downstream analysis. The MSI and IMC data was then integrated at the MSI pixel level, where for each MSI pixel the cell type proportion according to area and the mean IMC marker intensity was calculated.

#### Association testing

Testing for glycan-celltype association at the bulk level was done by calculating, for each ROI, the glycan mean intensity followed by fitting a linear regression of bulk glycan intensity vs. sex, TMA and sample_type (one of: No Tumor, Adjacent to Tumor, Tumor). Testing at the domain aggregated level was done accordingly by aggregation per ROI, glycan and domain, followed by linear modeling. Statistical analysis to determine associations between cell types and glycans at the MSI pixel-level was done using the two-level approach described in MIMIC [9]. First for each ROI a simultaneous autoregressive model is fit with the glycans as response and the (per-pixel normalized) cell type areas as covariates. Second, the estimated parameters are used in a linear regression model (also incorporating other covariates to account for technical artifacts) to obtain mean associations across samples, as well as condition specific changes of association.

## Results and Discussion

### Feasibility and design constraints of same-section IMC–AAXL–MALDI integration

We first evaluated the feasibility of combining IMC and AAXL–MALDI MSI of N-glycans could be performed on the same FFPE section and, if so, in which order. Our workflow combined on-tissue sialic-acid derivatization for isomer-sensitive N-glycan MSI [18] with a validated IMC antibody panel (Figure 1A). For each tissue core, the MALDI MSI acquisition area encompassed the full IMC region of interest (Figure 1B). Optimal laser power for this study was determined emperically through testing a range of laser powers S1. Although IMC involves laser ablation, optimization of laser power (–1 dBa) produced minimal visible tissue disturbance (Figure 1C). Cell types were identified through normalized and median marker expression per cluster were visualized using heatmaps, violin plots, and statistical summaries (Figures S2-S3).

**Figure 1.**
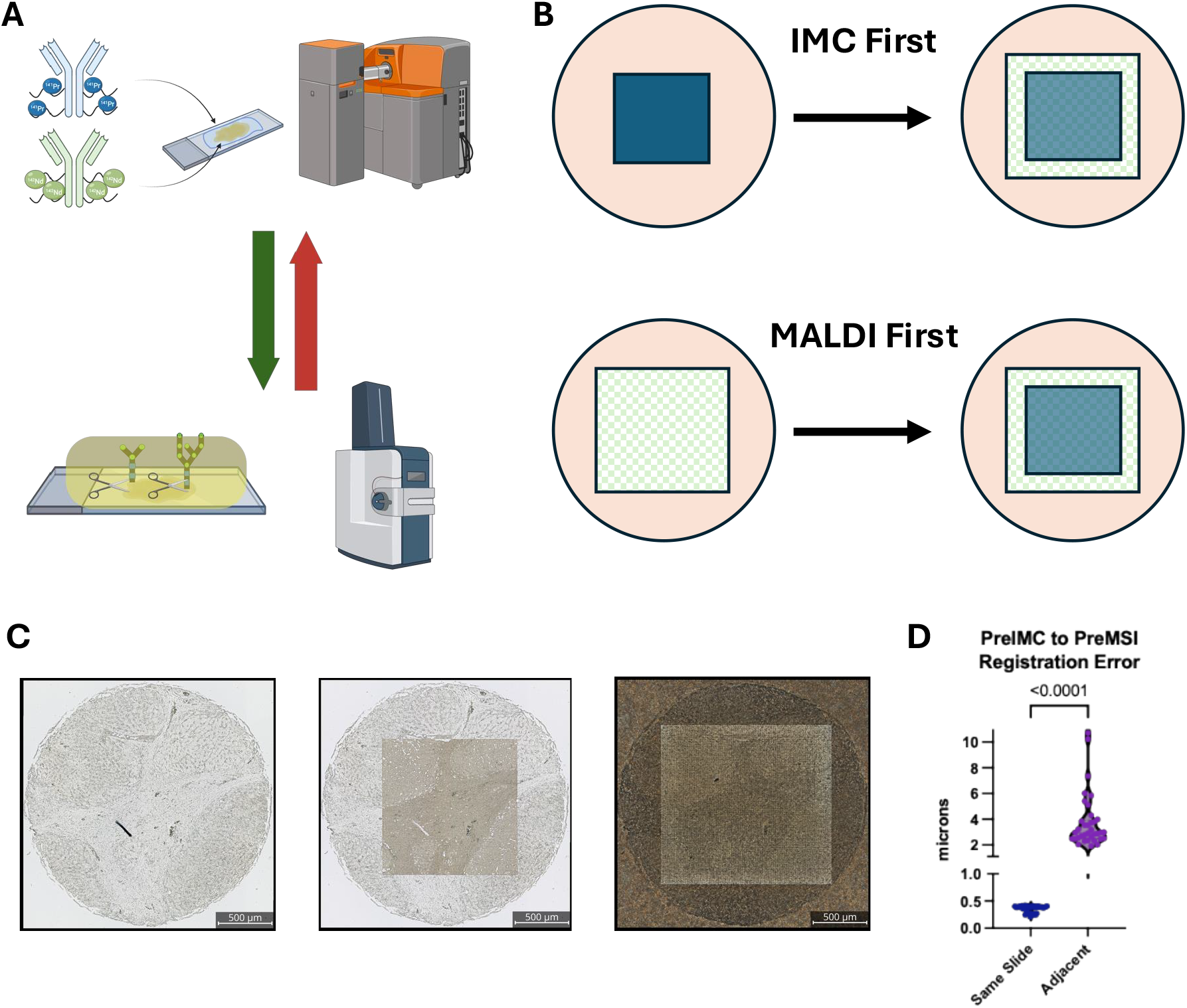
Sequential IMC–AAXL–MALDI workflow and rationale for same-section acquisition. **A**. Overview of IMC staining with a metal-tagged antibody panel and AAXL–MALDI MSI for isomer-sensitive N-glycan imaging. **B**. Two acquisition orders were compared: IMC followed by AAXL–MALDI (IMC-first) or AAXL–MALDI followed by IMC (MALDI-first). **C**. Representative tissue appearance before acquisition, after IMC ablation, and after AAXL–MALDI. Optimized IMC laser power (–1 dBa) produced minimal visible tissue disturbance. **D**. Comparison of registration errors between same-section and adjacent-section IMC–MALDI acquisition. n=36, paired t-test.

To assess acquisition strategies, we compared adjacent-section versus same-section acquisition. Same-section imaging required less tissue, minimized variation in cell-type composition between samples, and substantially reduced registration error (Figure 1D and Figure S4). As shown previously [9], registration imprecision diminishes statistical power to detect celltype-glycan associations. Given the consistently lower registration error on the same slide, all subsequent analyses focused on same-section IMC and AAXL–MALDI MSI. This decision established a stringent test of modality compatibility while maximizing analytical resolution for downstream multimodal integration.

### Chemical incompatibility of AAXL–MALDI-first processing with IMC antibody integrity

We then evaluated how AAXL–MALDI first processing affects IMC antibody performance. When IMC was performed prior to AAXL–MALDI (“IMC-first”), staining was strong and specific across epithelial, immune, and nuclear markers (Figure 2A). In contrast, AAXL–MALDI-first processing resulted in a near-complete loss of detectable signal for the same markers (Figure 2B).

**Figure 2.**
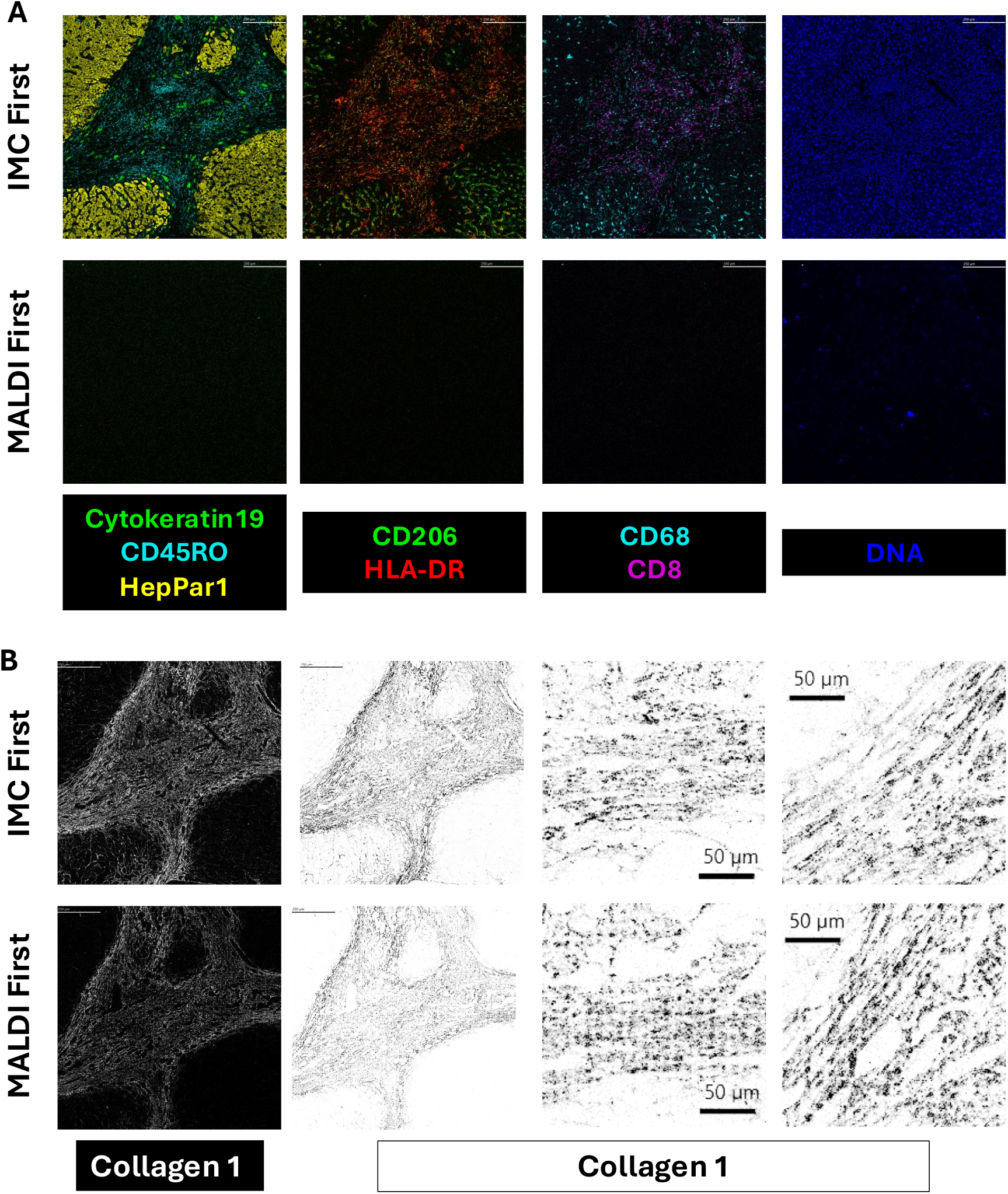
AAXL–MALDI-first processing disrupts IMC antibody staining. **A**. Pseudocolor IMC images showing staining for epithelial (CK19, HepPar1), immune (CD45RO, CD68, CD206, HLA-DR, CD8), and nuclear (DNA) markers when IMC is performed before AAXL–MALDI (IMC-first, top row). or after AAXL–MALDI (MALDI-first, bottom row). **B**. Collagen I staining in IMC first (top row) and AAXL–MALDI first (bottom row) highlighting the grid pattern corresponding to the MALDI laser footprint.

This disruption was global rather than epitope-specific and extended across antibody classes, indicating widespread loss of antibody-binding capacity rather than selective degradation. Importantly, this level of disruption was not observed in prior workflows using non-derivatized N-glycan MALDI MSI [13], indicating that the chemical steps required for AAXL sialic-acid stabilization—rather than laser ablation alone—are responsible for loss of antibody performance. The carbodiimide-mediated activation, nucleophilic amidation, elevated-temperature incubations, and repeated organic solvent washes required for sialic acid stabilization collectively impose a chemically aggressive environment that is incompatible with preserving antibody epitopes in FFPE tissue.

In the small subset of markers that remained detectable, staining intensity was substantially reduced relative to IMC-first (Figures S5-S6), rendering quantitative interpretation unreliable. These findings establish that AAXL–MALDI-first processing fundamentally compromises IMC antibody performance on the same tissue section and therefore precludes meaningful antibody-based imaging downstream.

### Preservation of AAXL–MALDI MSI following IMC-first acquisition

In contrast, we observed that performing IMC prior to AAXL–MALDI MSI preserved the analytical integrity of downstream glycan imaging. Spatial ion distributions for representative sialylated, fucosylated, and high mannose glycans (*m/z* 2137.766, 1905.634, 1809.639) were highly concordant between IMC-first and AAXL–MALDI-first conditions, with similar regional enrichment patterns across tissue domains (Figure 3A). Additional images of sialylated glycans showing concordance between the acquisition orders are shown in S7. Quantifying signal intensities within versus outside IMC ablation footprints revealed *<* 5% difference (Figure 3B), indicating minimal local laser-induced impact on subsequent glycan detection.

**Figure 3.**
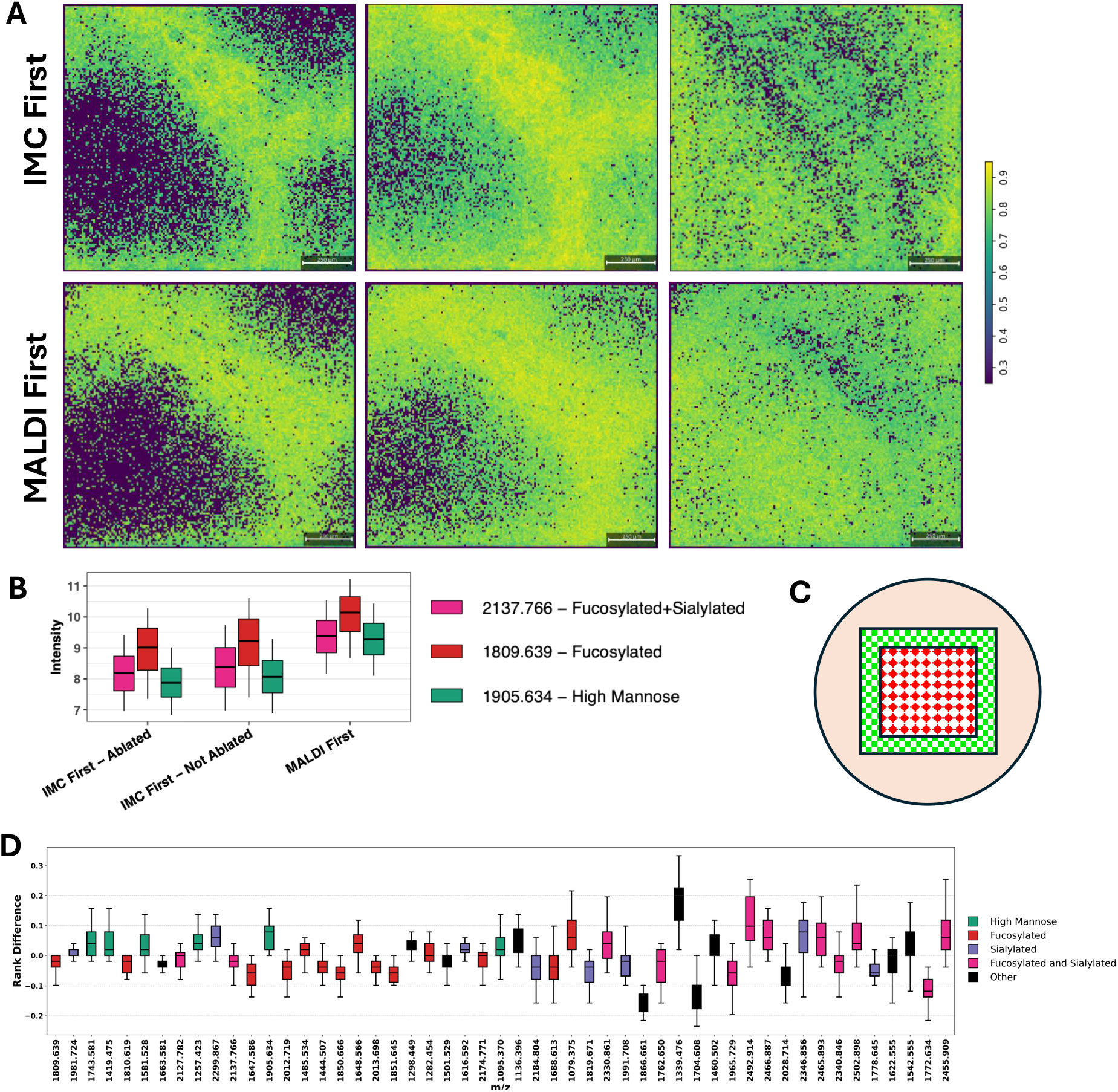
IMC-first preserves spatial and relative abundance patterns in AAXL–MALDI MSI. **A**. Ion images for representative N-glycans (*m/z* 2137.766, *α*2, 3 sialylated, Hex5dHex1HexNAc4NeuAc1; *m/z* 1809.639, fucosylated, Hex5dHex1HexNAc4; *m/z* 1905.634, high mannose, Hex9HexNAc2) comparing IMC-first and MALDI-first acquisition orders. **B**. Quantification of AAXL–MALDI intensities within versus outside the IMC ablation footprint. **C**. Graphical representation of regions quatified in B with IMC First - Ablated as red diamonds and IMC First - Not Ablated as green squares. **D**. Rank-order comparison of abundances of different glycans across regions of interest (ROIs).

A global attenuation in AAXL–MALDI signal ( ∼15%) was observed in IMC-first slides (Figure 3B); however, this shift was uniform across *m/z* species and comparable to standard batch-to-batch variation [24] [25]. Relative glycan abundance patterns were preserved: glycan rank-order analysis showed that > 90% of species changed < 10% in their ROI-level rank position between acquisition orders (Figure 3C).

Thus, while IMC-first acquisition introduces a mild and systematic reduction in absolute signal intensity, it does not distort spatial distributions, linkage-specific patterns, or relative abundance relationships. These properties are the primary determinants of biological interpretability in AAXL–MALDI MSI and confirm that IMC-first sequencing is analytically compatible with isomer-resolved glycan imaging.

### Robust multimodal integration at domain and single-cell resolution following IMC-first sequencing

Finally, we assessed whether IMC-first sequencing preserves the multimodal relationships required for integrated analysis. Using the MIMIC co-registration pipeline, domain-level associations between glycans and major histological features (e.g., hepatocyte-rich versus collagen-rich regions) were highly concordant across acquisition orders, with similar direction and magnitude of enrichment patterns (Figure 4A). Directionality and magnitude of enrichment patterns were highly consistent, indicating that spatial glycan–tissue relationships remain intact.

**Figure 4.**
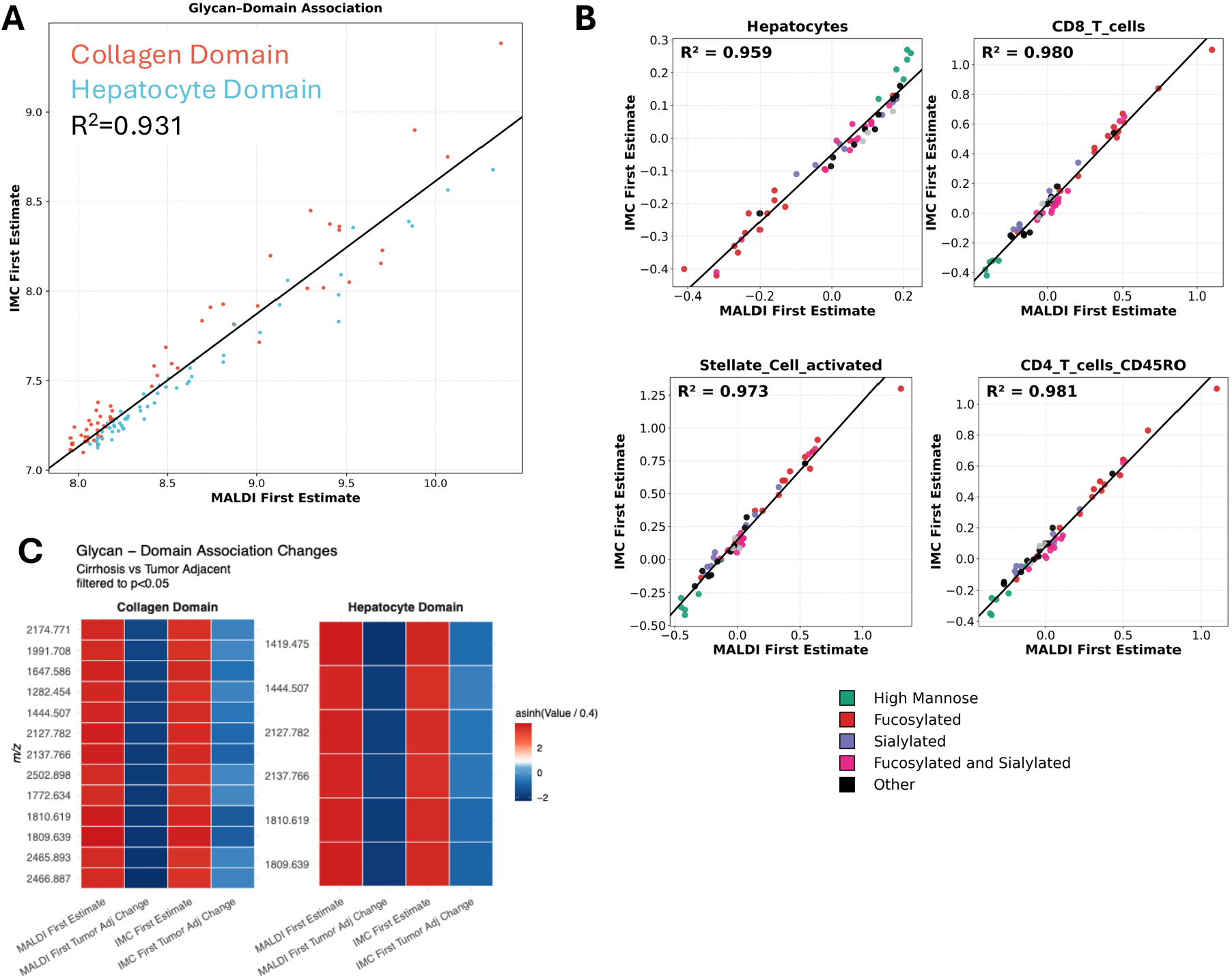
Domain- and single-cell–level glycan associations are preserved with IMC-first sequencing. Registration of the cell types annotated from the IMC-first slide to the MSI pixels from the same slide, and to the adjacent slide where MSI was performed first. Correlation of domain-level associations **A**. and cell-level association **B**. among glycan species between acquisition orders. **C**. Quantification of changes in glycan-domain association across sample type cirrhosis vs tumor adjacent.

At the single-cell level, correlations between AAXL–MALDI glycan features and IMC-resolved cell types—including hepatocytes, stellate cells, macrophages, and T-cell subsets—also remained strong (Figure 4B). Across cell populations, glycan–cell type associations showed high agreement between acquisition orders (median *R*^2^ ≈0.90). These results demonstrate that IMC-first sequencing maintains the domain-level and single-cell glycan–cell relationships necessary for integrated spatial sialylation analysis within the same FFPE section. We then assesed how glycan-domain associations change across biological conditions, in this case cirrhosis only (No Tumor) vs tumor-adjacent cirrhosis. We show in Figure 4C that both acquisition orders preserve the same biological story. There is a significant loss of glycosylation in tumor adjacent tissue compared to cirrhosis only. However, due to the attenuated signal on IMC First, this observation is also attenuated. Thus, signal attenuation as a result of IMC First may reduce the magnitude of observed change, but it does not affect the directionality. Collectively, these findings indicate that IMC-first acquisition not only preserves individual modality performance but also enables statistically robust, biologically interpretable cross-modal analyses within the same FFPE tissue section.

## Conclusions

This study establishes a clear and generalizable principle for multimodal spatial imaging: acquisition order is not a technical afterthought but a defining analytical parameter that determines whether integration is feasible or fundamentally compromised. For workflows combining antibody-based imaging with chemically intensive, derivatization-dependent mass spectrometry, physical tissue perturbation and chemical perturbation are not equivalent—and must be evaluated independently. Our data demonstrate that the chemically aggressive steps required for AAXL-based, isomer-resolved N-glycan imaging irreversibly disrupt antibody-binding capacity when applied prior to IMC, rendering same-section antibody imaging analytically invalid. In contrast, optimized IMC acquisition imposes only minimal physical perturbation and does not meaningfully distort downstream glycan spatial distributions, relative abundance structure, or multimodal associations. As a result, IMC-first sequencing uniquely satisfies the dual requirement of preserving antibody integrity while maintaining chemically faithful glycan MSI. Beyond defining a recommended acquisition order, this work provides a framework for rational multimodal workflow design. It highlights that compatibility cannot be inferred from prior lipid or non-derivatized glycan studies and must instead be evaluated in the context of the specific chemical transformations employed. These principles extend beyond sialylated N-glycan imaging and are broadly applicable to any spatial workflow combining antibody-based imaging with derivatization-dependent MSI chemistries. By establishing a validated, same-section IMC-first AAXL–MALDI workflow, this study enables high-resolution integration of cellular phenotype with linkage- and isomer-specific glycan biology in FFPE tissues. More broadly, it underscores the necessity of chemistry-aware experimental design in next-generation multimodal spatial biology and provides a practical blueprint for integrating complex molecular readouts without sacrificing analytical rigor.

## References

(1) Buchberger, A. R.; DeLaney, K.; Johnson, J.; Li, L. Analytical Chemistry 2018, 90, 240–265.

(2) Lin, J.-R.; Fallahi-Sichani, M.; Chen, J.-Y.; Sorger, P. K. Current Protocols in Chemical Biology 2016, 8, 251–264.

(3) Black, S.; Phillips, D.; Hickey, J. W.; Kennedy-Darling, J.; Venkataraaman, V. G.; Samusik, N.; Goltsev, Y.; Schürch, C. M.; Nolan, G. P. Nature Protocols 2021, 16, 3802–3835.

(4) Angelo, M.; Bendall, S. C.; Finck, R.; Hale, M. B.; Hitzman, C.; Borowsky, A. D.; Levenson, R. M.; Lowe, J. B.; Liu, S. D.; Zhao, S.; Natkunam, Y.; Nolan, G. P. Nature Medicine 2014, 20, 436–442.

(5) Giesen, C.; Wang, H. A. O.; Schapiro, D.; Zivanovic, N.; Jacobs, A.; Hattendorf, B.; Schüffler, P. J.; Grolimund, D.; Buhmann, J. M.; Brandt, S.; Varga, Z.; Wild, P. J.; Günther, D.; Bodenmiller, B. Nature Methods 2014, 11, 417–422.

(6) Bollhagen, A.; Whipman, J.; Coelho, R.; Heinzelmann-Schwarz, V.; Jacob, F.; Bodenmiller, B. Nature Methods 2025, 22, 2601–2608.

(7) Zhu, X.; Xu, T.; Peng, C.; Wu, S. Frontiers in Chemistry 2022, Volume 9 -2021.

(8) Grgic, A.; Cuypers, E.; Dubois, L. J.; Ellis, S. R.; Heeren, R. M. A. Journal of Proteome Research 2024, 23, 5372–5379.

(9) Gerber, R.; Griner, J.; Guglietta, S.; Krieg, C.; Robinson, M. D. bioRxiv 2025, 2025.07.08.663623.

(10) Esselman, A. B.; Moser, F. A.; Tideman, L. E.; Migas, L. G.; Djambazova, K. V.; Colley, M. E.; Pingry, E. L.; Patterson, N. H.; Farrow, M. A.; Yang, H.; Fogo, A. B.; de Caestecker, M.; Van de Plas, R.; Spraggins, J. M. Kidney International 2025, 107, 332–337.

(11) Nunes, J. B.; Ijsselsteijn, M. E.; Abdelaal, T.; Ursem, R.; van der Ploeg, M.; Giera, M.; Everts, B.; Mahfouz, A.; Heijs, B.; de Miranda, N. F. C. C. Nature Methods 2024, 21, 1796–1800.

(12) Veličković, D.; Purkerson, J.; Bhotika, H.; Huyck, H.; Clair, G.; Pryhuber, G. S.; Anderton, C. Molecular Omics 2025, 21, 334–342.

(13) Dunne, J.; Griner, J.; Romeo, M.; Macdonald, J.; Krieg, C.; Lim, M.; Yagnik, G.; Rothschild, K. J.; Drake, R. R.; Mehta, A. S.; Angel, P. M. Analytical and Bioanalytical Chemistry 2023, 415, 7011– 7024.

(14) Wu, R.-Q. et al. Immunity 2023, 56, 180–192.e11.

(15) Balneger, N.; Cornelissen, L. A. M.; Wassink, M.; Moons, S. J.; Boltje, T. J.; Bar-Ephraim, Y. E.; Das, K. K.; Søndergaard, J. N.; Büll, C.; Adema, G. J. Cellular and Molecular Life Sciences 2022, 79, 98.

(16) Weber, K. S. C.; Alon, R.; Klickstein, L. B. Inflammation 2004, 28, 177–188.

(17) Wheeler, S. F.; Domann, P.; Harvey, D. J. Rapid Communications in Mass Spectrometry 2009, 23, 303–312.

(18) Lu, X.; McDowell, C. T.; Blaschke, C. R. K.; Liu, L.; Grimsley, G.; Wisniewski, L.; Gao, C.; Mehta, S.; Haab, B. B.; Angel, P. M.; Drake, R. R. Analytical Chemistry 2023, 95, 7475–7486.

(19) Windhager, J.; Bodenmiller, B.; Eling, N. bioRxiv 2021, 2021.11.12.468357.

(20) Greenwald, N. F. et al. Nature Biotechnology 2022, 40, 555–565.

(21) Korsunsky, I.; Millard, N.; Fan, J.; Slowikowski, K.; Zhang, F.; Wei, K.; Baglaenko, Y.; Brenner, M.; Loh, P.-r.; Raychaudhuri, S. Nature Methods 2019, 16, 1289–1296.

(22) Lun, A.; McCarthy, D.; Marioni, J. F1000Research 2016, 5, DOI: 10.12688/f1000research.9501.2.

(23) Bemis, K. A.; Föll, M. C.; Guo, D.; Lakkimsetty, S. S.; Vitek, O. Nature Methods 2023, 20, 1883–1886.

(24) Balluff, B.; Hopf, C.; Porta Siegel, T.; Grabsch, H. I.; Heeren, R. M. A. Journal of the American Society for Mass Spectrometry 2021, 32, 628–635.

(25) Huang, L.; Kim, Y.; Balluff, B.; Cillero-Pastor, B. Analytical Chemistry 2025, 97, 10919–10928.

