## Supplementary figures and methods for "Resolving Sialylated N-Glycans and Immune Cell Landscapes Using a Unified Same-Section IMC–MSI Workflow"

### Supplementary Tables

**Table S1**. Clinical characteristics of TMA cohort

| **Characteristic** | **n (%) / mean [95% CI]** |
|---|---|
| **Sex** | |
| Female | 8 (33.3%) |
| Male | 16 (66.7%) |
| **Cancer** | |
| Non-cancer | 8 (33.3%) |
| Cancer | 16 (66.7%) |
| **Labs** | |
| Age | 61.9 [58.9–65.0] |
| BMI | 33.3 [30.1–36.6] |
| AST | 56.9 [47.7–66.2] |
| ALT | 39.6 [31.9–47.2] |
| ALK Phos | 164.8 [118.7–211.0] |
| Bilirubin | 4.4 [0.7–8.0] |
| Platelet | 98.9 [72.7–125.1] |
| Total Protein | 6.8 [6.4–7.2] |
| Albumin | 3.0 [2.8–3.3] |

**Table S2**. Imaging Mass Cytometry Antibodies

| Isotope | Metal | Marker | Vendor | RRID | Clone | Cell Type |
|---|---|---|---|---|---|---|
| 141 | Pr | aSMA | Standard BioTools | 2890139 | 1A4 | Activated fibroblasts, myofibroblasts, smooth muscle cells |
| 143 | Nd | Vimentin | Standard BioTools | 3106905 | D21H3 | Mesenchymal cells, fibroblasts, endothelial cells |
| 144 | Nd | CD14 | Standard BioTools | 2924314 | EPR3653 | Monocytes, macrophages |
| 145 | Nd | CD31 | Standard BioTools | 3086679 | EPR3094 | Endothelial cells |
| 146 | Nd | CD16 | Standard BioTools | 3106906 | EPR16784 | NK cells, neutrophils, macrophages |
| 148 | Nd | Pan-keratin | Standard BioTools | 2938626 | C11 | All epithelial cells |
| 149 | Sm | CD11b | Standard BioTools | 2891189 | EPR1344 | Myeloid cells (monocytes, macrophages, neutrophils, dendritic cells) |
| 150 | Nd | CK19 | Biolegend | 439773 | A53-B/A2 | Cholangiocytes |
| 151 | Eu | CD163 | Standard BioTools | 3106930 | EDHu-1 | M2 macrophages |
| 152 | Sm | CD45 | Standard BioTools | 2909538 | D9M8I | All leukocytes |
| 153 | Eu | CD206 | Standard BioTools | 3106931 | 5C11 | M2 macrophages |
| 155 | Gd | FoxP3 | Standard BioTools | 3106908 | PCH101 | Regulatory T cells (Tregs) |
| 156 | Gd | CD4 | Standard BioTools | 3094536 | EPR6855 | CD4+ T cells |
| 158 | Gd | E-cadherin | Standard BioTools | 2893074 | 24E10 | Epithelial cells, hepatocytes |
| 159 | Tb | CD68 | Standard BioTools | 2810859 | KP1 | Macrophages |
| 160 | Gd | CD66b | Standard BioTools | 3106932 | BLR111H | Neutrophils |
| 161 | Dy | CD20 | Standard BioTools | 2811016 | H1 | B cells |
| 162 | Dy | CD8a | Standard BioTools | 2661802 | CD8/144B | CD8+ T cells |
| 164 | Dy | CD32b | Abcam | 726388 | EP887Y | Liver Sinusoidal Endothelial Cells (LSECs) |
| 167 | Er | Granzyme B | Standard BioTools | 2811057 | EPR20129-217 | Cytotoxic effector function |
| 168 | Er | Ki-67 | Standard BioTools | 2810856 | B56 | Proliferating cells |
| 169 | Tm | Collagen 1 | Standard BioTools | 2810857 | Polyclonal | ECM, fibroblasts, hepatic stellate cells, myofibroblasts |
| 170 | Er | CD3 | Standard BioTools | 2661807 | Polyclonal | T cells |
| 171 | Yb | Histone H3 | Standard BioTools | 3662096 | D1H2 | All nucleated cells |
| 172 | Yb | Beta Catenin | Standard BioTools | 3662097 | 5H10 | Epithelial cells, endothelial cells, hepatocytes |
| 173 | Yb | CD45RO | Standard BioTools | 2811052 | UCHL1 | Memory T cells |
| 174 | Yb | HLA-DR | Standard BioTools | 2924389 | LN3 | Antigen-presenting cells (APCs), activated T cells |
| 175 | Lu | HepPar1 | Santa Cruz | 781327 | OCH1E5 | Hepatocytes |
| 176 | Yb | CCR6 | Standard BioTools | 3662098 | G034E3 | Memory T cells, Th17 cells, dendritic cells |
| 195 | Pt | Seg 1 | Standard BioTools | 3662095 | NA | Segmentation marker |
| 196 | Pt | Seg 2 | Standard BioTools | 3662095 | NA | Segmentation marker |
| 198 | Pt | Seg 3 | Standard BioTools | 3662095 | NA | Segmentation marker |

**Table S3**. MALDI Acquisition Parameters.

| | |
|---|---|
| **Transfer** | |
| MALDI Plate Offset | 70.0 V |
| Deflection 1 Delta | 90.0 V |
| Funnel 1 RF | 500.0 Vpp |
| isCID Energy | 8.0 eV |
| Funnel 2 RF | 500.0 Vpp |
| Multipole RF | 500.0 Vpp |
| **Collision Cell** | |
| Collision Energy | 15.0 eV |
| Collision RF | 4000.0 Vpp |
| **Quadrupole** | |
| Ion Energy | 18.0 eV |
| Low Mass | 600.00 m/z |
| **Focus Pre TOF** | |
| Transfer Time | 120.0 $\mu$s |
| Pre Pulse Storage | 30 $\mu$s |
| **Laser Settings** | |
| Shots per Pixel | 200 |

#### Supplementary Figures

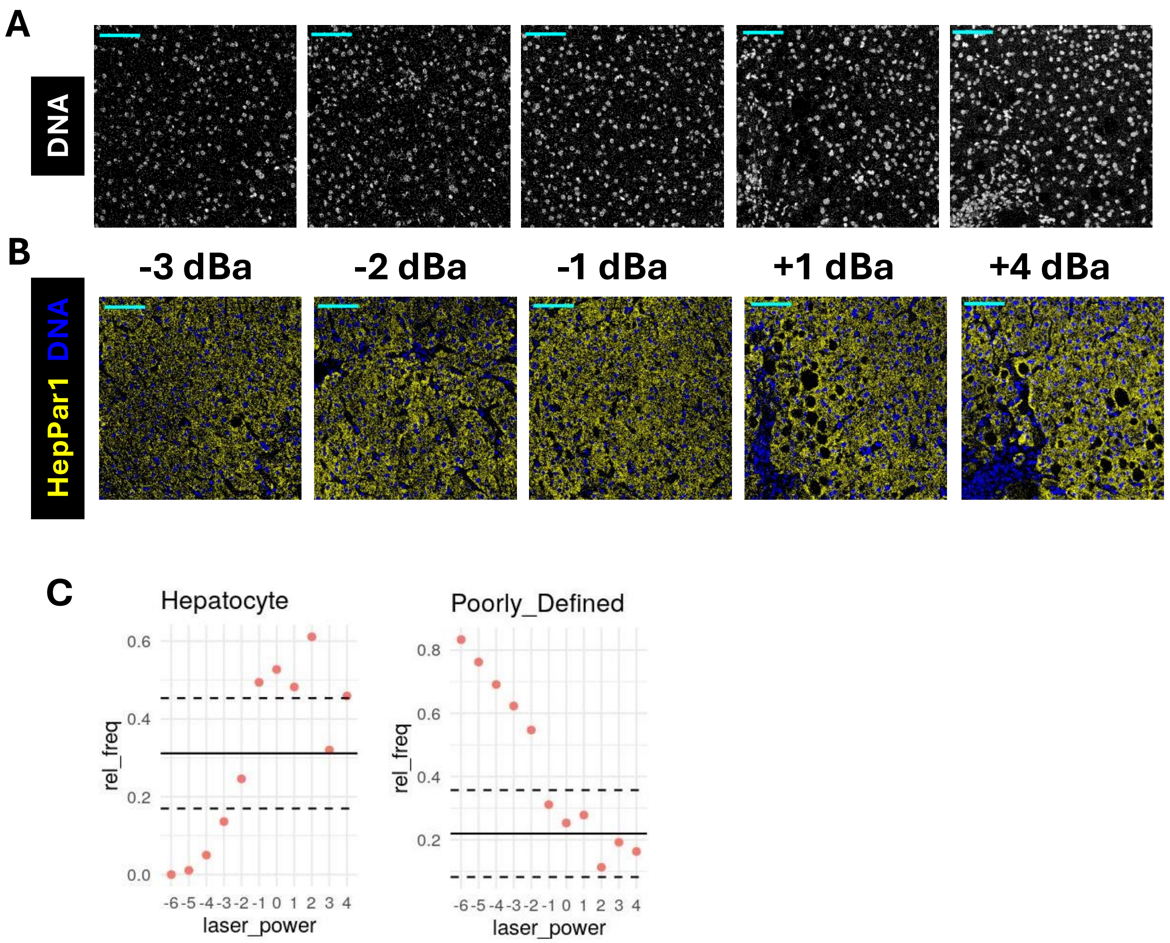


**Figure S1**. **Empirical optimization of laser power for imaging mass cytometry acquisition.** Representative images of DNA intercalator signal (A) and HepPar1 staining (B) across a range of laser power settings. (C) Quantification of cell annotation outcomes across laser power settings.

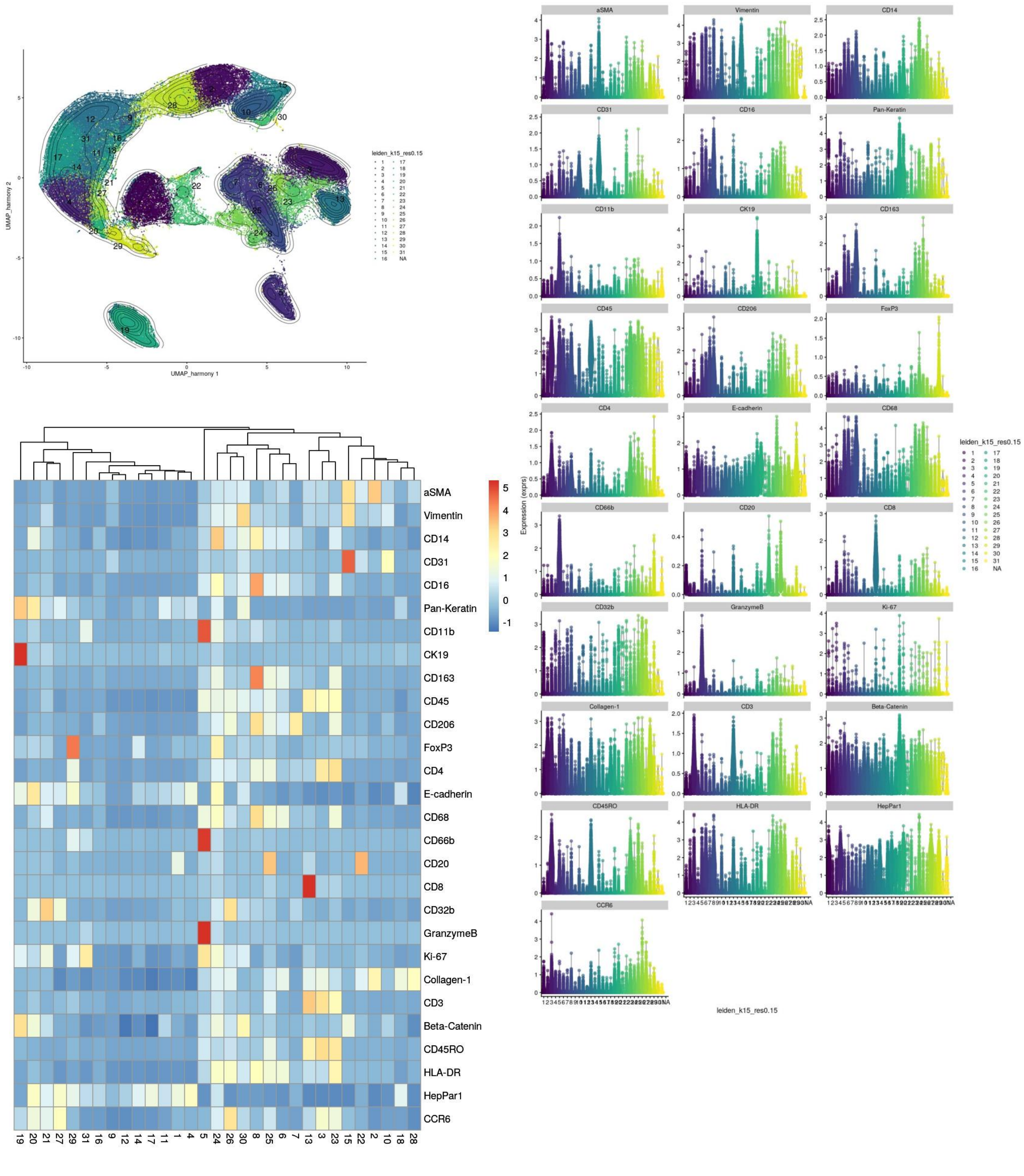


**Figure S2**. **Dimensionality reduction and unsupervised clustering prior to manual cluster consolidation.** Cells were embedded using UMAP following Leiden community detection (k = 20 nearest neighbors, resolution = 0.15). UMAP projections display cluster assignments, with accompanying heatmaps and violin plots showing median marker expression per cluster.

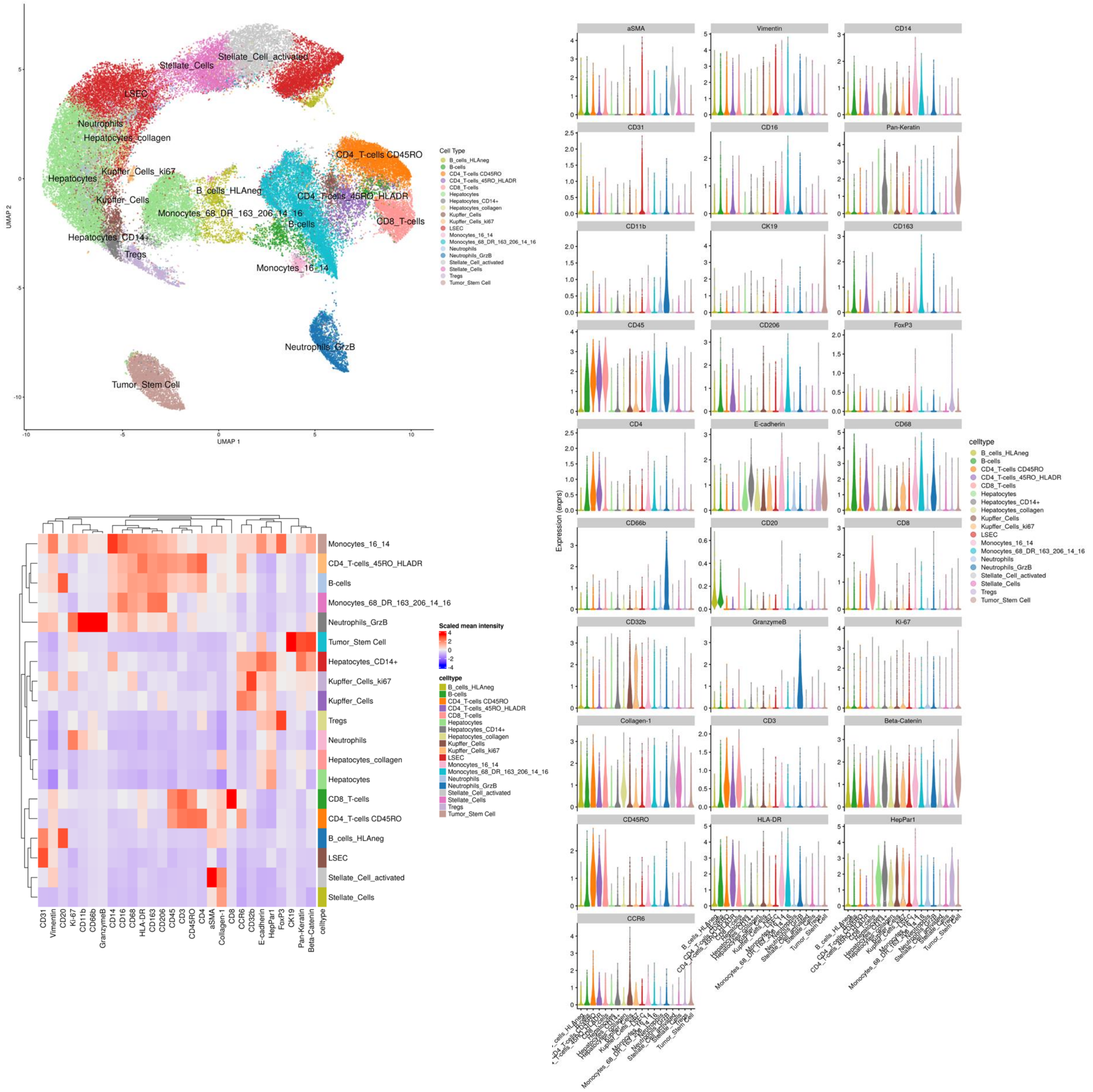


**Figure S3**. **Dimensionality reduction and cell-type annotation following cluster consolidation.** The UMAP projection displays the final cell-type labels. The heatmap summarizes scaled median marker intensity across annotated cell types, and accompanying violin plots show marker expression distributions within each annotated population.

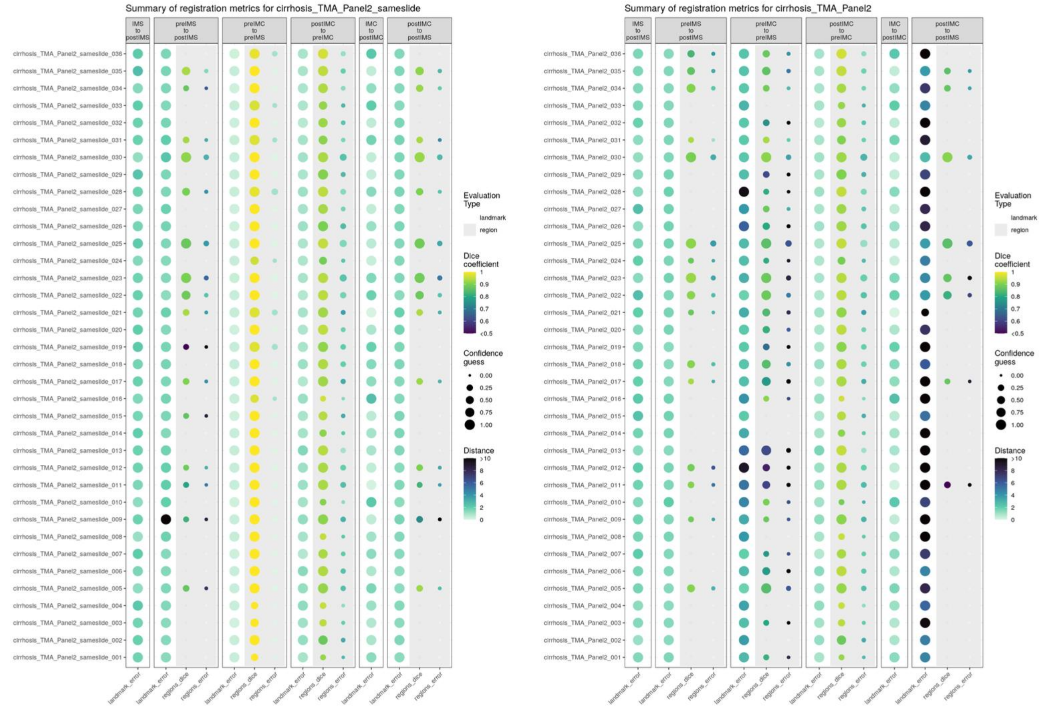


**Figure S4**. **Summary of image registration metrics across TMA cores.** Each row represents an individual core, and columns correspond to pairwise registration steps (IMS to postIMS, preIMS to postIMS, preIMC to preIMS, postIMC to preIMC, IMC to postIMC, and postIMC to postIMS). Point color encodes Dice similarity coefficient, and an additional color scale denotes landmark distance error where applicable. The left panel displays same-slide (IMC First) registration performance, while the right panel displays adjacent-slide (MALDI First) registration performance.

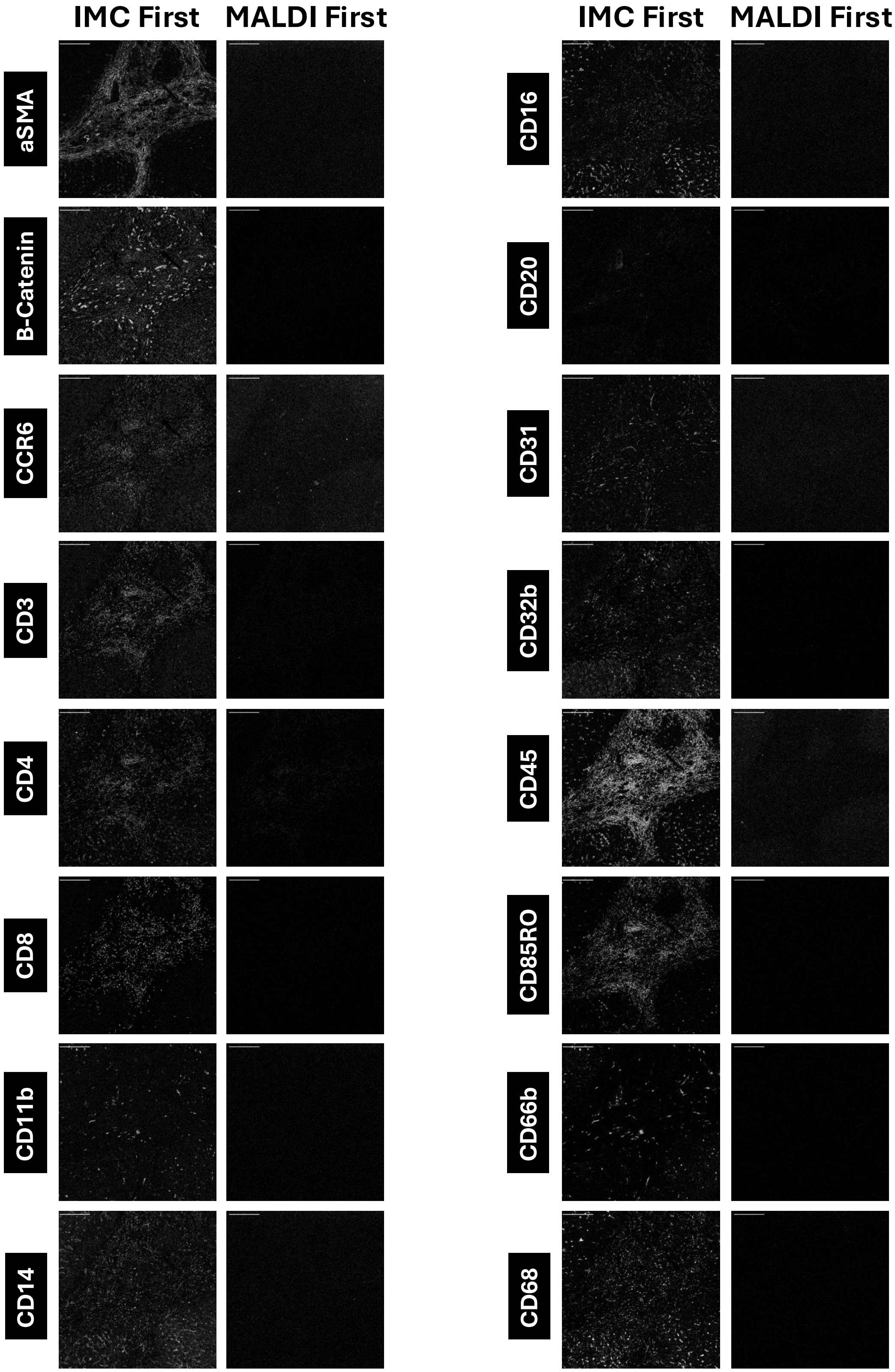


**Figure S5**. **Representative single-channel IMC images comparing acquisition order.** Grayscale intensity images are displayed with IMC performed prior to MALDI (IMC First) and with MALDI prior to IMC (MALDI First). All images are shown at identical spatial scale and contrast settings.

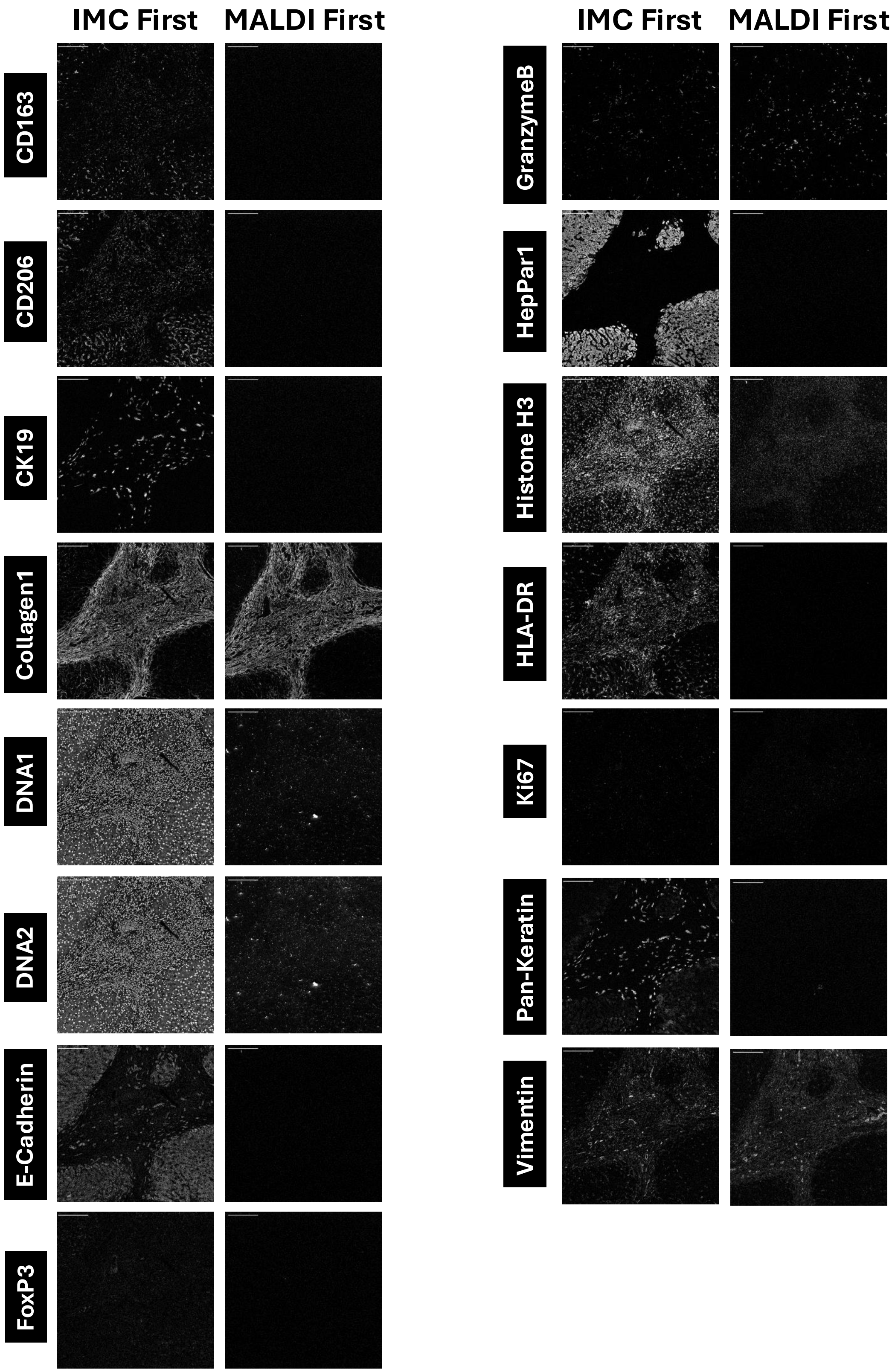


**Figure S6**. **Additional single-channel IMC images comparing acquisition order.** Grayscale TIFF exports are shown for each marker under IMC-first and MALDI-first conditions.

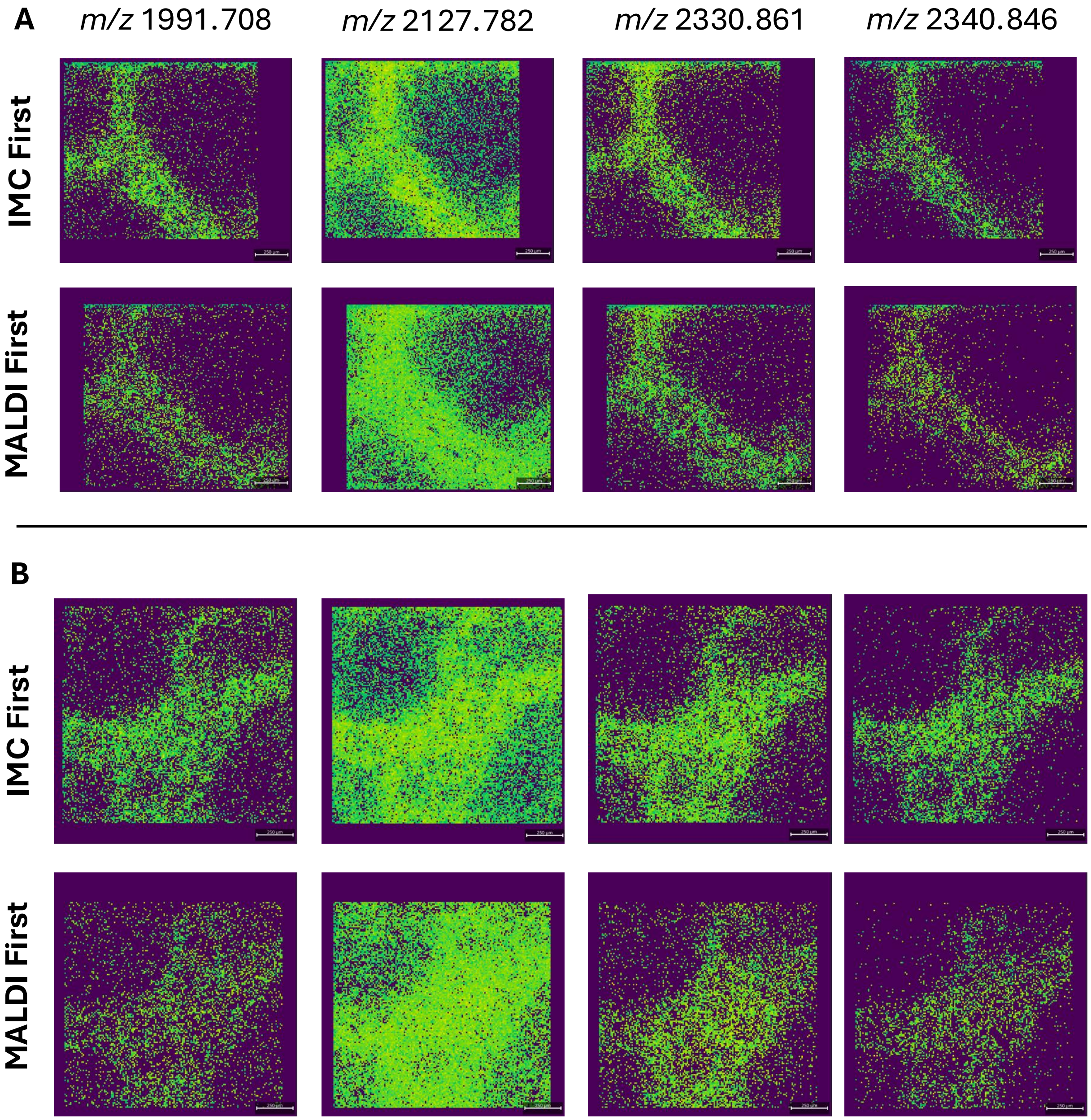


**Figure S7**. **Additional sialylated glycan ion images.** Left to right: $m/z$ 1991.708 ($Hex_5HexNAc_4NeuAc_1$, isomer $\alpha 2,3$); $m/z$ 2127.782 ($Hex_5dHex_1HexNAc_4NeuAc_1$, isomer $\alpha 2,6$); $m/z$ 2330.861 ($Hex_5dHex_1HexNAc_5NeuAc_1$, isomer $\alpha 2,6$); and $m/z$ 2340.846 ($Hex_5dHex_1HexNAc_5NeuAc_1$, isomer $\alpha 2,3$). Panel A shows representative core C1; Panel B shows representative core C2.

### Integrated IMC–AAXL–PNGase F Workflow (MIMIC)

#### Scope

This protocol describes an integrated workflow in which FFPE tissue sections are:

1. stained for Imaging Mass Cytometry (IMC),
2. derivatized using the AAXL method for differential labeling of sialylated glycans, and
3. processed for Matrix-Assisted Laser Desorption/Ionization (MALDI) mass spectrometry imaging (MSI) via on-tissue PNGase F digestion and matrix application.

All steps are performed sequentially on the same slide.

#### Definitions

**Imaging Mass Cytometry (IMC).** Imaging Mass Cytometry detects spatial distributions of lanthanide-labelled antibodies at near–single-cell resolution.

**AAXL derivatization.** AAXL derivatization stabilizes and distinguishes $\alpha 2,3$ vs. $\alpha 2,6$ sialylated glycans via differential amidation and lactone formation. It produces the following characteristic mass shifts:

| | |
|---|---|
| $\alpha 2,3$ sialic acids | +37.0316 Da |
| $\alpha 2,6$ sialic acids | +27.0473 Da |

**PNGase F digestion.** PNGase F digestion releases N-linked glycans for MALDI-MS imaging.

### Reagents and Equipment

#### Equipment

Coplin jars; 5-slide mailers; 60 °C oven; antigen retrieval devices (Decloaking Chamber™ NxGen (BioCare Medical)); humidity chambers; vacuum desiccator; flatbed document scanner; 40x brigtfield whole slide scanner (Ex. Hamamatsu NanoZoomer); HTX TM-Sprayer; syringe pump; glass incubation chamber; glass coverslip.

#### Reagents

**IMC Reagents** Xylenes; 200-proof ethanol; 95%/70% ethanol; HPLC water; Tris-EDTA pH 9 buffer; TBS with Triton X-100; BSA; PBS; PBS with Triton; Intercalator-Ir.

**AAXL Reagents** EDC (1-ethyl-3-(3-dimethylaminopropyl)carbodiimide); HOBt (1-hydroxybenzotriazole); dimethylamine; propargylamine; DMSO; ethanol; chloroform; glacial acetic acid; trifluoroacetic acid (TFA); HPLC water.

**PNGase F / MALDI Reagents** Citraconic anhydride buffer; 12 M HCl; PNGase F; CHCA ($\alpha$-cyano-4-hydroxycinnamic acid); trifluoroacetic acid (TFA); HPLC water; acetonitrile.

### Procedure

#### IMC Workflow

##### Tissue Sectioning

1. Section FFPE tissue at 5 $\mu$m onto positively charged slides.
2. Ensure tissue is centered with at least 2 mm clearance from slide edges to allow IMC ablation.

**Dewaxing and Rehydration**

1. Bake slides at 60 °C for 1 h.
2. Dewax slides in xylenes (3 min ×2).
3. Rehydrate sequentially:
   (a) 100% ethanol (1 min ×2),
   (b) 95% ethanol (1 min),
   (c) 70% ethanol (1 min).
4. Rinse in HPLC water (3 min ×2).

**IMC Antigen Retrieval (Tris–EDTA pH 9)**

1. Dilute Tris–EDTA pH 9 buffer 1:100 in HPLC water.
2. Load slides into a 5-slide mailer; fully cover tissue with buffer.
3. Place mailer into decloaking chamber.
4. Ensure the slide mailer is **mostly but not fully submerged** in the chamber water bath.
5. Run Program 4 (20 min at 95 °C; total cycle approx. 30 min).
6. Cool gradually via three water-exchange steps (5 min each).
7. Finish with a final rinse in 100% HPLC water.

**IMC Blocking**

1. Encircle tissue completely with PAP pen; allow boundary to dry.
2. Prepare blocking buffer: TBS + 0.1% Triton X-100 + 3% BSA.
3. Apply blocking buffer within PAP boundary.
4. Incubate for 1 h at room temperature in a humidity chamber.

**IMC Staining and Washes**

1. Prepare antibody master mix in TBS + 0.1% Triton X-100 + 1% BSA.
2. Apply antibody mix and incubate overnight at 4 °C in a humidity chamber.
3. Wash slides:
   (a) PBS + 0.2% Triton (5 min ×2),
   (b) PBS (5 min ×2).
4. Apply Intercalator-Ir (1:1000) for 30 min at room temperature.
5. Wash in HPLC water (5 min ×2).
6. Dry slides in a desiccator for $\geq$ 5 min before scanning.

**Scan 1 — Pre-IMC Brightfield Slide Scan (40×)**
Performed after IMC staining, prior to IMC acquisition. Required for alignment and downstream co-registration in the MIMIC pipeline.

##### IMC Laser Optimization and Ablation

1. Load slide into Hyperion per SOP.
2. Optimize laser power empirically to achieve:
   (a) Adequate signal intensity for low-abundance markers.
   (b) Minimal visible tissue ablation.
3. Acquire IMC data across all ROIs.

**Scan 2 — Post-IMC Brightfield Slide Scan (40×)**
Performed after IMC ablation. Captures ablation zone and used to link IMC signal and cell masks to on-tissue location.

---

#### AAXL Derivatization (Post-IMC)

##### Reagent Preparation

**1st Amidation Solution** Dissolve 42.2 mg HOBt in 500 $\mu$L DMSO; add 15.8 $\mu$L dimethylamine; add 22 $\mu$L EDC. Mix and use immediately.

**Wash Solutions** Carnoy's solution (60% ethanol / 30% chloroform / 10% acetic acid); 0.1% TFA in ethanol.

##### First Amidation Reaction

1. Apply 200–400 $\mu$L of the 1st reaction solution to tissue.
2. Cover with glass coverslip, avoiding bubbles.
3. Incubate for 1 h at 60 °C.
4. Remove coverslip and wash with 200 $\mu$L DMSO (×3).

##### Second Amidation Reaction

1. Prepare 2nd Amidation solution by mixing 300 $\mu$L propargylamine with 700 $\mu$L DMSO. Prepare fresh.
2. Apply 200–400 $\mu$L of the 2nd Amidation solution.
3. Apply a clean coverslip, avoiding bubbles.
4. Incubate for 2 h at 60 °C.

##### Washing

1. Remove coverslip and place in glass coplin jar.
2. Wash in 100% ethanol (2 min ×2).
3. Wash in Carnoy's solution (10 min ×2).
4. Wash in 100% ethanol (2 min ×2).
5. Pour over 0.1% TFA in ethanol.

6. Rinse in HPLC water (3 min ×2).
7. Dry slides in desiccator (5 min to overnight).

---

#### pH 3 Citraconic Antigen Retrieval (Pre-PNGase F)

##### Buffer Preparation

1. Combine 25 mL distilled water + 25 $\mu$L citraconic anhydride + 2 $\mu$L 12 M HCl.
2. Bring final volume to 50 mL; verify pH $\approx$ 3.

##### Heat-Induced Antigen Retrieval (Decloaking Chamber, Program 4)

1. Load slides into a 5-slide mailer and fully cover tissue with citraconic buffer.
2. Place mailer into decloaking chamber.
3. Ensure the slide mailer is **mostly but not fully submerged** in the chamber water bath.
4. Run Program 4 (20 min at 95 °C).
5. Cool using three water-exchange steps (5 min each).
6. Final rinse in distilled water; dry in desiccator for 5 min.

---

#### On-Tissue PNGase F Digest

##### PNGase F Preparation

Prepare PNGase F at 0.1 $\mu$g/$\mu$L in HPLC-grade water.

##### Spraying PNGase F

1. Load PNGase F solution into TM-Sprayer syringe; remove bubbles.
2. Spray using:
   - Nozzle 45 °C
   - 15 passes
   - Velocity 1200
   - 3 mm spacing
   - 10 psi nitrogen
   - Syringe pump 25 $\mu$L/min

##### Enzymatic Incubation

1. Prepare humidified incubation chamber (water-saturated towel + supports).
2. Pre-warm to $38.5 \pm 1.5\,^\circ$C for $\geq 30$ min.
3. Incubate for 2 h at $38.5 \pm 1.5\,^\circ$C.

---

#### Matrix Application

##### CHCA Preparation

Dissolve CHCA at 7 mg/mL in 50% acetonitrile + 0.1% TFA. Sonicate 5 min; filter through 0.2 $\mu$m.

##### Spraying CHCA

1. Load matrix into sprayer syringe.
2. Spray using:
   - Nozzle 80 °C
   - 10 passes
   - Velocity 1300
   - 2.5 mm spacing
   - 10 psi nitrogen
   - Pump rate 0.1 mL/min
3. Store slides in a desiccator.

**Scan 3 — Post-MALDI MSI Brightfield Slide Scan (40×)**
Performed after MALDI MSI acquisition. Used in the MIMIC workflow to link spectra to their on-tissue location.
